# Sound perception in realistic surgery scenarios: Towards EEG-based auditory work strain measures for medical personnel

**DOI:** 10.1101/2024.05.07.592873

**Authors:** Marc Rosenkranz, Thorge Haupt, Manuela Jaeger, Verena N. Uslar, Martin G. Bleichner

**Affiliations:** Neurophysiology of Everyday Life Group, Department of Psychology, Carl von Ossietzky Universität Oldenburg, Oldenburg, Germany; Pius-Hospital Oldenburg, University Hospital for Visceral Surgery, Carl von Ossietzky Universität Oldenburg, Oldenburg, Germany; Research Center for Neurosensory Science, Carl von Ossietzky Universität Oldenburg, Oldenburg, Germany

## Abstract

Surgical personnel face various stressors in the workplace, including environmental sounds. Mobile electroencephalography (EEG) offers a promising approach for objectively measuring how individuals perceive sounds. In this study, we utilized mobile EEG to explore how a realistic soundscape is perceived during simulated laparoscopic surgery. To examine the varying demands placed on personnel in different situations, we manipulated the cognitive demand during the surgical task, using a memory task. To assess responses to the soundscape, we calculated event-related potentials for distinct sound events and temporal response functions for the ongoing soundscape. Although participants reported varying degrees of demand under different conditions, no significant effects were observed on surgical task performance or EEG parameters. However, changes in surgical task performance and EEG parameters over time were noted, while subjective results remained consistent over time. These findings highlight the importance of using multiple measures to fully understand the complex relationship between sound processing and cognitive demand. Furthermore, in the context of combined EEG and audio recordings in real-life scenarios, sparse representations of the soundscape have advantages over more detailed representations. Our results indicate that both types of representations are equally effective in eliciting neural responses. Overall, this study marks a significant step towards objectively investigating sound processing in applied settings.

## Introduction

Surgical personnel often experience high levels of stress, which can lead to severe health problems such as burnout^1^ or hypertension^2,3^. One cause of stress is distractions due to the environment of the operating room (OR)^4,5^. The soundscape in the OR is highly complex^6^, comprising of sounds that are crucial to the surgery (e.g, communication, tool usage), as well as sounds that are not crucial to the surgery (e.g., instrument clattering, phone ringing) and could be minimized to improve the work environment^7^. The accumulation of the different sounds sources, leads to high sound levels which are often perceived as distracting and stressful^8–13^. To guide interventions that aim at reducing stress induced by auditory distractions it is important to understand and measure how ongoing sounds in the OR affect the personnel.

An objective evaluation of the subjectively experienced burden of the OR soundscape is challenging. Previous studies that have focused on the effect of sound on performance (i.e., surgery task performance) have found that only under extreme and unnatural conditions, such as dichotic listening to two pieces of music^14,15^, but not under more natural conditions, such as a single stream of OR sounds^16–21^, more mistakes in the surgery task were observable. However, medical personnel may strive to perform at high levels under adverse working conditions because mistakes can have serious consequences for the patient. Thus, the work strain surgeons experience can not necessarily be inferred from the surgery task performance. Therefore, additional measures are required to objectively assess the auditory strain in the OR.

To measure sound perception in the OR objectively, electroencephalography (EEG) is a promising method. Given its temporal resolution, it is particularly useful for measuring responses that are time-locked to sounds. By analyzing event-related potentials (ERPs) the neural response to individual sound events can be examined, revealing perceptual and cognitive processes such as whether a sound is considered task-relevant^22^. Another approach involves the use of temporal response functions (TRFs) to study responses to continuous sound streams^23^, which expands the use of EEG to naturalistic soundscapes. ERPs and TRFs can be measured outside of the laboratory while individuals are freely moving^24,25^ and to complex soundscapes while an audio-visual-motor task is being performed^26^. The development of head-mounted EEG systems and wireless data transmission allows for unrestricted mobility for the wearer, making it applicable in various work environments^27^ and enabling measurements over extended periods of time^28^.

To leverage EEG in the OR and investigate auditory strain, we examined neural responses to continuous soundscapes in OR-related situations. In the current study, we utilized a laparoscopic simulator that required bi-manual control, and incorporated a soundscape representative of typical OR environments^29^, providing a realistic and challenging environment for participants. As most OR sounds are irrelevant to surgeons^30^, we presented a task-irrelevant soundscape including, for example, ventilation humming, tool clattering, and tool usage. This element of realism ensures that our findings are applicable to actual surgical settings. Additionally, it provides insights into the relationship between common OR sounds and cognitive processing.

In the complex environment of surgical operations, understanding the impact of varying cognitive demand on performance and sound distraction is crucial for both patient safety and the efficiency of medical procedures^7,12^. Traditional studies varied the demand by employing dual-task methodologies, where participants are required to engage in a primary task while also responding to secondary stimuli, to study the interference of additional cognitive demands on task performance^31–35^. However, such approaches may not accurately reflect the nuanced cognitive challenges inherent to surgical tasks, where the demand is often internalized and involves the simultaneous management of multiple streams, such as concentrating on the procedure while blending out distractions, without overt responses. Recognizing this gap, our study proposes a novel approach to simulate the cognitive demands placed on surgeons by incorporating a passive memory task. This methodology is designed to simulate the maintenance and manipulation of information without the use of an overt task response. By focusing on this aspect, our study aims to provide a realistic assessment of how internal cognitive processes impact surgical performance and sound perception. We investigated demand-dependent changes in EEG responses to the soundscape, as well as subjective and behavioural measures. We focused on ERP and TRF time-windows typically found for sounds, namely the N1, P2, and N2 time-window^36–38^. Previous studies reported mixed findings regarding the effect of varying demand on these time-windows^39^. Therefore, we investigated the effect of varying demand on each time-window.

Surgeons are frequently required to perform complex procedures for extended periods. This naturally leads to fluctuations in cognitive demand and how the environment is perceived. For example, sounds that were once easily ignored can become sources of distraction, or vice versa^12^. This change in sensory processing, compounded by varying demand, underscores the critical need to consider the temporal dynamics during the performance of surgical tasks. Therefore, we explored changes over time for the ERP and TRF time-windows in response to the soundscape, as well as for the subjective and behavioral measures. At last, there are practical considerations when opting to use TRFs in an applied context. Since the derivation of TRFs require information of the soundscape, sound recordings have to adhere to privacy concerns. To test whether representations of the soundscape void of personal information, produce similar results as rich representations we derived sound onset marker^40^ in addition to the commonly used acoustic envelope^23^. Sound onsets only indicate sound occurrence, providing a data protected way of sound recording. Previous research has demonstrated that onsets and envelopes yield comparable results for computing TRFs^41,42^. To replicate this finding and strengthen the applicability of TRFs in real-life settings we compared the TRFs computed from the onsets and the envelopes of the soundscape.

The objective of this research was twofold. Firstly, it aimed to examine the influence of a realistic task under varying levels of cognitive demand on the individual processing of task-irrelevant auditory stimuli, reflecting an OR soundscape. This investigation integrated both subjective assessments (i.e., self-reports) and objective metrics (i.e., neural responses to irrelevant sounds and behavioral performance) to provide a comprehensive understanding of sound and distractor processing in work environments. Secondly, the study investigated the processing of continuous versus discrete acoustic features, offering insights into the response of the auditory system to different aspect of the soundscape.

## Methods

This study involving human participants was approved by the Medizinische Ethikkommission, Carl von Ossietzky Universität Oldenburg, Oldenburg (2021-031) and performed in accordance with the Declaration of Helsinki. The participants provided their written informed consent to participate in this study. This study was preregistered after data collection https://doi.org/10.17605/OSF.IO/AE3UY. All changes to the preregistration can be found in the supplementary material.

### Participants

23 medical students were recruited through an online announcement on the university board and word-of-mouth (age range: 19-24; 16 female). One participant was excluded due to data loss during recording resulting in 22 included participants. We based the sample size on our previous studies in which we measured reliable ERPs and TRFs during a complex and dynamic task^26^. Four participants had previous experience with a laparoscopic simulator and 17 attended at least once a surgery as an observer. All participants received monetary reimbursement. Eligibility criteria included: enrollment as a medical student and self-reported normal hearing, normal or corrected vision, no psychological or neurological condition, and right-handedness.

### Paradigm

Figure 1 illustrates the setup and paradigm. Participants performed a laparoscopic surgery task while presented with a continuous soundscape. To vary the cognitive demand during the surgery task, we adapted a serial recall paradigm: Prior to the surgery task, participants had to remember a list of letters, and received the instruction to silently repeat those letters while performing the surgery task. After the surgery task, they should recall the letters.

**Figure 1.**
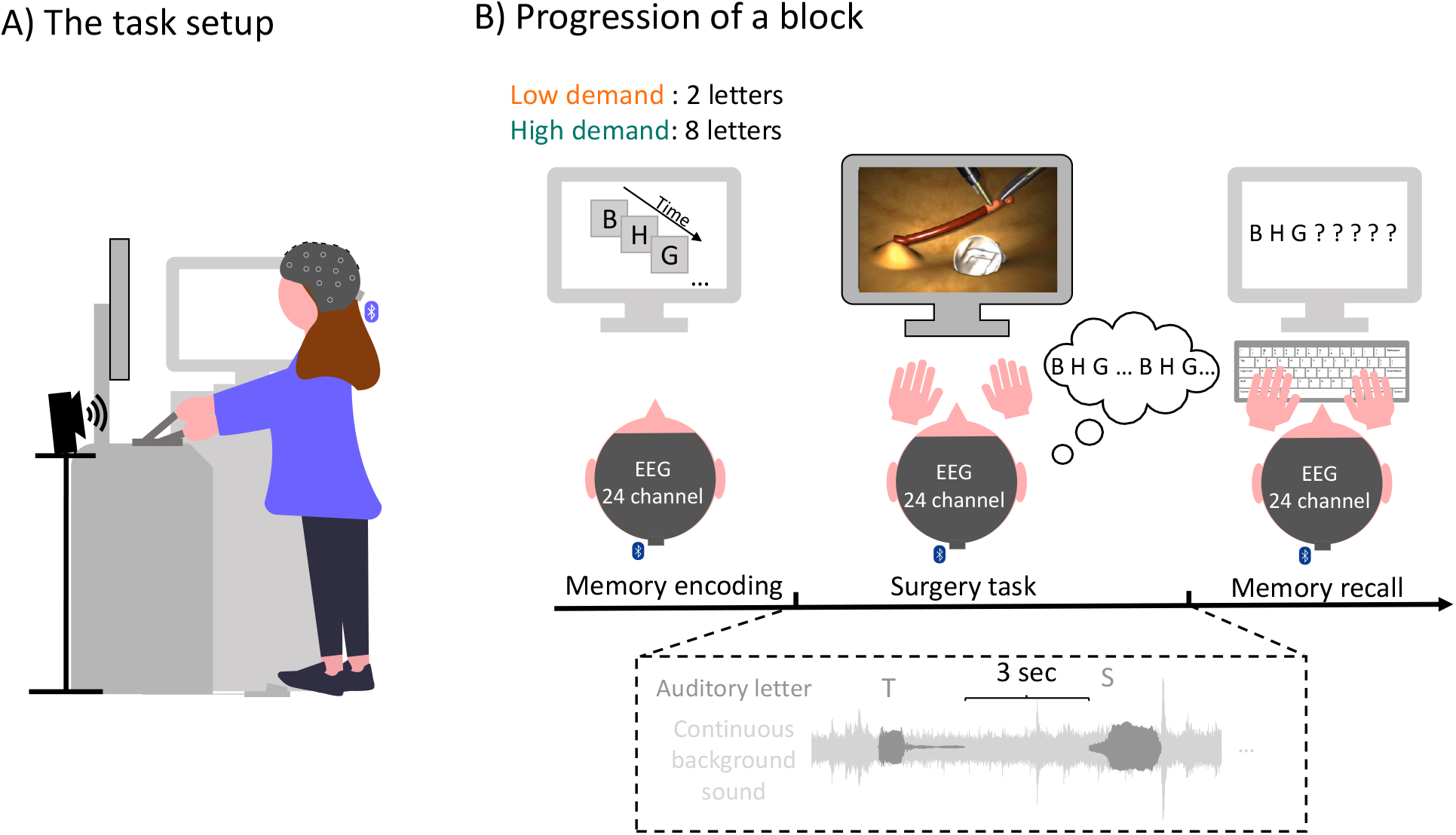
A) Participants were standing throughout the entire experiment either in front of the surgery simulator or the screen for the memory task which were positioned in 90° to each other. The speaker were positioned at chest height and participants were equipped with a mobile EEG cap. B) At the start of the block participant were presented with either two (low-demand condition) or eight (high-demand condition) letters which they had to remember. During the surgery they were presented with a continuous realistic soundscape of an operation room and with overlayed spoken letters. They were instructed to silently repeat the letters during the surgery task. After finishing the surgery task they should enter the letters and fill out a workload questionnaire. An experimental block ended with feedback regarding their memory performance.

The soundscape contained two parts. Firstly, to mimic the actual surgical environment, the playback of a continuous OR soundscape was presented. Secondly, to increase the distractive potential of the soundscape, we presented spoken letters, which are commonly used as distractors in classic serial recall paradigms^43,44^. Participants did not receive any instruction on what to do with the soundscape, and were free to disregard it.

After retrieval of the letters, participants rated their perceived demand during the task and received feedback regarding their memory performance. Each participant performed a total of 28 experimental blocks, 14 of each condition, in randomized order. During the entire experiment, participants were standing either in front of the screen for the memory task (Samsung, SyncMaster P2470) or in front of the surgery simulator. The monitor and simulator were position in 90° to each other. Further details can be found below.

#### Surgery task

For the surgery task, the LabSim^®^ (Surgical Science, Sweden) simulator was used. This simulator is used to train hand-eye coordination for laparoscopic surgeries and provides realistic graphics and tactile feedback. The setup includes specialized instruments for both hands, a foot pedal for additional controls, and a screen that displays the virtual surgical instruments in action.

The specific task which the participant performed throughout the experiment is called “cutting”. In this task, participants needed to grasp a vessel with the right-hand instrument, stretch it, and cut it with an electrical cutting device which was handled with the left hand. To activate the cutter, a foot paddle must be pressed. The vessel must then be placed in a small bag. The task ends when three parts of the vessel are removed.

Participants were informed that their performance would be evaluated based on the duration to complete the surgery task, mistakes, and tissue damage. Mistakes included rupturing the vessel (e.g., by exerting too much pull on the vessel) or dropping the cut vessel, thus a block could have zero to three mistakes. Touching the tissue with the instrument caused immediate visual feedback as the screen received a red shade. After each task, they received visual feedback about their overall performance.

#### Memory task

A serial recall task was adapted to vary the cognitive demand during the surgery task. Although the serial recall task is a classic working memory task, we refrain from calling it one, as the retention interval (i.e., the performance of the surgery task) was too long. Prior to the surgery task, participants were asked to remember the visually presented letters in the correct order. For the low-demand condition two letters and for the high-demand condition eight letters had to be remembered. The letters were randomly selected from a set of twelve letters (B,C,D,F,H,K,L,M,P,Q,S,T) without replacement. Letters were presented in black on a gray screen for 1000 ms each with an inter-stimulus-interval of 500 ms.

After the surgery task, either two or eight question marks indicated that the letters should be entered using a keyboard. Participants could enter an “X” for letters they could not remember and could correct themselves.

#### Soundscape

The OR playback was recorded using a field recorder which was positioned close to the surgery table during a visceral surgery at the University Hospital Oldenburg^29^. The recording contains a variety of sounds, such as ventilation noise, beeps from monitoring devices, instrument clatter, and instrument sounds. Intelligible speech was removed after the recording for privacy reasons, however, unintelligible muttering and non-vocal sounds such as coughing were preserved. The recording lasts approximately 1 hour. In the first block, the recording starts at a random time point and continuous chronologically for every subsequent block. If the recording reached the end, it started at the beginning. The soundscape started automatically after the letters of the memory task were presented. To prevent clicking noise when the audio starts the starting 500 milliseconds were faded in.

In addition to the realistic soundscape we presented a sequence of individual spoken letters. For the spoken letters, four letters were drawn from the same set of letters as the memory task, but never coincided with the to-be-remembered letters of a block. To ensure that the letters were presented equally often within a block, the four letters were presented as consecutive groups but were randomized within a group. The same letter was never presented consecutively. The inter-trial-interval between letters was three seconds. Letters were generated using a text-to-speech program (Notevibes, accessed 2021) and spoken by the same female voice. The letters were clearly audible in the soundscape recording and had an-on and offset ramp of ten milliseconds. All sounds (i.e., the recording and letters) were sampled at a rate of 48 kHz and presented to the participant using Psychtoolbox 3 for MATLAB^45^ (v3.0.17), a t.amp E4-130 amplifier (Thomann GmbH, Burgebrach, Germany) and presented as a stereo signal using two iLoudMTM loudspeakers (IK Multimedia Production srl, Modena, Italy). The loudspeakers were vertically tilted upwards (20°) and located in front of the participant to the right and to the left of the LabSim^®^ at chest height. The distance between loudspeakers was 0.5 m and the distance between loudspeaker and ear was 1.2 m. The sound pressure level was 45-55 dB SPL, measured at the place of the participants head.

#### Subjective workload assessment

To assess the subjectively perceived demand during each block we included three workload related questions. For a good representation of our research question we chose two items from the NASA-TLX^46^, namely effort (“How hard did you have to work to accomplish your level of performance?”) and frustration (“How insecure, discouraged, irritated, stressed, and annoyed were you?”) and one question from the SURG-TLX^47^, namely distraction (“How distracting was the operating environment”). Each question was answered on a visual analog scale ranging from 0 to 20^46,47^.

#### Training

To familiarize themselves with the tasks, participants engaged in a series of practice blocks. During these sessions, the experimenter remained nearby, providing guidance and support to help participants navigate through the tasks. First, they performed the memory task twice, once with two letters and once with eight letters. Second, they performed a basic surgery task (i.e., instrument navigation) without the memory task or sounds to familiarize themselves with the LabSim^®^. Third, they performed the cutting task without the memory task or sounds. Lastly, participants performed two training blocks that were identical with the blocks of the main experiment, first for the low-demand condition and then for the high-demand condition. After the training, the experiment started and participants performed the experimental blocks on their own.

### Data acquisition

Participants were asked to wash their hair on the day of the recording and to not use hair styling products. EEG data were recorded using a wireless 24-channel amplifier (SMARTING, mBrainTrain, Belgrade, Serbia) attached to the back of the EEG cap (EasyCap GmbH, Hersching, Germany) with Ag/AgCl passive electrodes at 10-20 layout positions (Fp1 Fp2 AFz Fz F3 F4 F7 F8 Cz C3 C4 T7 T8 CPz Pz M1 M2 P3 P4 P7 P8 POz O1 O2) with the reference and ground electrode at position FCz and Fpz, respectively. The data were recorded using a sampling rate of 500 Hz, and transmitted via Bluetooth from the amplifier to a Bluetooth dongle (BlueSoleil) that was plugged into a computer.

After fitting the cap, the skin was cleaned using 70% alcohol. Abrasive gel (Abralyt HiCl, Easycap GmbH, Germany) was used for reducing electrical impedance and ensuring high-quality signal. Impedances were kept below 20 kΩ at the beginning and again checked at the end of the recording using the SMARTING Streamer software (v3.4.3; mBrainTrain, Belgrade, Serbia). ECG data were also recorded on a laptop but were not part of the current analyses.

Experimental markers (e.g., sound markers) were generated using the lab streaming layer library^48^ (v1.14). The ECG recording laptop and EEG recording computer were connected via Lan. To synchronize all data streams, EEG data, ECG data, and experiment marker were collected in the Lab Recorder software (v1.14, https://github.com/labstreaminglayer/App-LabRecorder) and saved as one .xdf file on the EEG recording computer.

### Preprocessing

**EEG** The EEG data were analyzed using EEGLAB^49^ (v2022.0) in MATLAB R2020b (The MathWorks, Natick, MA, United States). For each participant, the continuous data were filtered with Hamming windowed FIR filter using the EEGLAB default settings: (1) high-pass: passband edge = 0.5 Hz (filter order = 3300, transition bandwidth = 0.5 Hz, cutoff frequency (−6dB) = 0.25 Hz); (2) low-pass: passband edge = 30 Hz (filter order = 220, transition bandwidth = 7.5 Hz, cutoff frequency (−6dB) = 33.75 Hz). For further analysis, only the EEG data during sound presentation were included, i.e., data during memory encoding and retrieval were removed. To minimize artifacts from switching between the surgery task screen and the memory task screen we removed the first and last five seconds of EEG data during sound presentation. Bad EEG channels were automatically rejected using the EEGLAB function *clean_artifacts* from the *clean raw data* plugin^50^ following a procedure described by Klug & Gramann (2021)^51^. The function was executed over ten iterations with the following parameters: ChannelCriterion (0.8); ChannelMaxBrokenTime (0.5). The remaining parameters were turned off. If a channel was rejected in at least 50% of iterations, it was removed. A maximum of 5 channels could be removed to ensure that enough data were available for channel reconstruction. On average 0.72 (*±* 1.28) channels were rejected.

After bad channel removal, the data were cleaned from artifacts using infomax independent component analysis (ICA). For ICA, a copy of the preprocessed data were created and high-pass filtered (passband edge = 1 Hz, filter order = 825, transition bandwidth = 1 Hz, cutoff frequency (−6dB) = 0.5 Hz), and cut into consecutive epochs of one second. Improbable epochs with a global (all channels) or local (single channel) threshold exceeding 5 standard deviations were automatically rejected using the *jointprob* function. ICA decomposition was applied to the remaining epochs. The resulting components were back-projected on the original preprocessed, but uncleaned data. The components were then classified using the EEGLAB toolbox *ICLabel*^52^ with the ‘lite’ classifier which is better at detecting muscle artifacts than the default classifier^53^. Components belonging to the categories eye blink and movement or muscle movement with 60% confidence were removed. Note, that the ICLabel classifier did not classify all components correctly because it was trained on stationary data with a larger electrode setup than ours. Therefore, we manually checked the components and made the following adjustments: We detected ICs located at the mastoids, probably from muscle movement (see Supplementary Figure S2 for an example). As the mastoids were used for re-referencing we manually removed these. Furthermore, lateral eye movement also required manual removal in some cases. Afterwards, previously rejected channels were interpolated using spherical interpolation. Lastly, channels were re-referenced to the linked mastoids (M1/M2).

#### Audio

For technical reasons, a constant delay between the marker that indicates a sound onset and the actual sound presentation was quantified beforehand. To correct for this constant delay the marker was shifted by 30 ms. Furthermore, the onsets of the auditory letters were also corrected. The letters were embedded in a constant sound stream which might lead to an energetic masking effect of the first few milliseconds of a letter. This affects when a letter becomes audible and therefore, the time when a brain response occurs. To obtain a better estimate when the participants could hear the letters, we used the OnsetDetector app^40^ implemented in MATLAB which determined the first energetic peak of the letters. The markers were shifted between 0 to 12.83 ms (Supplementary figure S1).

In order to relate the ongoing soundscape to the ongoing neural response, acoustic features were extracted. For this, we only used the OR playback (i.e., without the letters). From the OR playback we extracted and compared three feature vectors, namely the envelope of the raw OR playback, the envelope of the noise-reduced OR playback, and the onsets of the OR playback.

The envelope of the OR playback was extracted using the *mTRFenvelope* function^23,54^ with default inputs. To reduce the noise in the OR playback from ventilation and running machines, we used a Wiener filter implemented in MATLAB^55,56^. For this, we first high-pass filtered the soundscape at 1 Hz (filter order = 1000, transition bandwidth = 0.5 Hz, cutoff frequency (−6dB) = 0.00004 Hz). We then estimated the power spectral density of the noise using the first second of the OR playback, as it was representative of the static noise in the OR playback. The noise estimate was then subtracted from the remaining signal. Afterwards, we extracted the envelope from the noise-reduced OR playback using the *mTRFenvelope* function.

Onsets were calculated using the OnsetDetector App^40^ implemented in MATLAB. As we aim to detect onsets in naturalistic settings the raw audio was used. The resulting feature vector contained zeros (i.e., no onset) and ones (i.e., onsets).

### EEG analysis

We performed two types of analysis. An ERP analysis to study the event-related responses to the onset of the acoustic letters, and a TRF based analysis to study the response to the ongoing OR playback.

#### ERP calculation

ERP analysis was performed for the auditory letters. For each letter, epochs from -200 to 600 ms with respect to the stimulus onset were extracted. A baseline correction from -200 to 0 ms prior to stimulus onset was performed. Improbable epochs with a global (all channels) or local (single channel) threshold exceeding 3 standard deviations were automatically rejected using the *jointprob* function. We then computed the average response of each participant and block.

#### TRF calculation

A forward modeling approach was used to compute a temporal response function (TRF) that characterizes the brain’s temporal response to a feature vector representing the auditory stimulus. To calculate the TRF we used the mTRF toolbox^23^. For the TRF analyses, EEG data were multiplied by factor .0313 for scaling (as suggested in the provided code by Crosse et al. (2016)^23^).

To evaluate which acoustic features best predicts the neural response we implemented a forward model based on individual EEG data using a 10-fold cross-validation approach. For this, we separated the blocks into 10 segments. With 28 blocks in total, each segment consisted of two to three successive blocks. We split the segments into training and testing data such that each segment was once test data and iterated through the following procedure. For the training procedure we determined a shrinkage regularization parameter *λ* using the *mTRFcrossval* function with a time range from 0 to 450 ms time lag and a lambda range from 10^*−*8^ to 10^8^. This resulted in a correlation value for each fold (i.e., number of TRF training blocks), lambda value, and channel. We averaged over folds and channels and used the lambda value that maximized the correlation for the subsequent TRF calculation. We then trained a forward model using the training blocks and the *mTRFtrain* function with time lags from 0 to 450 ms and the optimal lambda. The resulting model was used to predict the response of the test blocks using the *mTRFpredict* function. This resulted in a prediction value for each test block, channel, and iteration, which were averaged, leaving one prediction value per participant and acoustic feature.

Lastly, we computed a forward model for each block using *mTRFtrain* with time lags from -220 to 500 ms. For the individual optimal lambda value we chose the most frequently occurring one during the cross-validation procedure. As the scale of TRFs varied across participants, biasing statistical comparisons of amplitudes, we z-scored each participant’s TRF weights across time-points, channel, and blocks.

#### GED analysis

To evaluate amplitude differences of the ERP and TRF and avoid channel selection, we use generalized eigenvalue decomposition (GED) as a spatial filter following guidelines by Cohen (2022)^57^. In short, for each ERP and TRF time-window (i.e., N1/P2/N2) we computed a generic spatial filter across subjects that was then applied to the data. GED maximizes the contrast between a signal covariance matrix S and a reference covariance matrix R. For S, we computed N1, P2, and N2 time-windows and contrasted each time-window separately against the baseline period (i.e., -200 to 0 ms).

In detail, we first average ERPs and TRFs across blocks, resulting in one time-series for each participant. We then determined the N1/P2/N2 peak of each ERP and TRF. Regarding the ERPs, and TRFs calculated from the onsets we searched for the N1 peak between 80-150 ms, for the P2 peak between 150-250 ms, and for the N2 peak between 200-300 ms. The TRF calculated from the envelope showed earlier peaks, therefore we searched for the N1 peak between 50-120 ms, for the P2 peak between 120-220 ms, and for the N2 peak between 220-320 ms.

Second, we determined the channel with the largest amplitude for the N1/P2/N2 peak. Around the peak, we then calculated a time-window of *±*25 ms for the N1 and *±*50 ms for the P2/N2 peak.

Third, to obtain the corresponding GED filter weights we contrasted the ERP and TRF time-windows with the baseline period.

Specifically, we mean-centered the data of the time-windows and baseline period and computed the covariance matrices for either time periods. For each participant, the covariance matrix of interest (S) was computed for the time-windows and the reference covariance matrix (R) was computed for the baseline period.

We cleaned the covariance matrix S and R by first computing the average covariance matrix 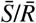 across participants. We then computed for each participant the Euclidean distance between the covariance S and 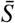, and R and 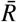. Covariance matrices which deviated from 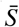 and 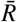 with more than 3 standard deviations were removed and 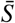 and 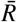 were computed again across the remaining covariance matrices. The 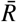 matrix was then regularized using a shrinkage regularization parameter of *λ* = 0.01. The resulting spatial filter maximizes the contrast between the time-windows and baseline period. For each ERP and TRF time-window, this resulted in a number of eigenvectors that is equal to the number of channels. The eigenvector with the largest eigenvalue should best separate the baseline and time-window. However, this does not guarantee physiological plausibility. We therefore investigated the GED components and reported when not using the GED component with the strongest eigenvalue. As a result we received a GED component time-series for each participant, ERP/TRF time-window, and block, and for the TRF additionally for each acoustic feature. Additionally, we computed a forward model for each component to investigate the physiological interpretability of the filter.

As eigenvectors are sign uncertain, we set appropriate signs for the GED components (negative for N1, N2, and positive for P2). Lastly, we averaged the amplitude of the GED component across the ERP and TRF time-windows to extract one amplitude value per time-window, participant, and block and for the TRF also per acoustic feature.

## Statistical analysis

All statistical analyses were performed in RStudio (v. 2021.09.0).

Prediction values of the acoustic feature were compared using a Wilcoxon signed rank test.

We computed regression models for the subjective workload questions, behavioral responses, and GED component mean amplitudes for the ERP and TRF time-windows (i.e., N1/P2/N2). For these analyses, we started with a model including ‘participant’ as a random intercept and ‘condition’ as a fixed effect. ‘Condition’ contained two categories, i.e., low- and high-demand which were coded 0 and 1, respectively. We explored time-on-task effects (i.e., they were not part of the preregistration) by adding the block number as a continuous predictor ‘time’ in a second model. In the third and most complex model an interaction term between condition and time was added:

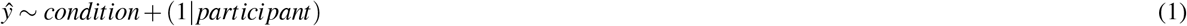

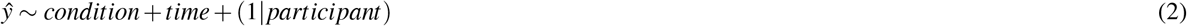

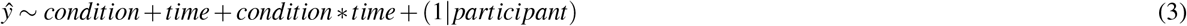

To test whether the models improved by adding predictors we used likelihood ratio testing. If the model fit did not improve, results from the first model were reported.

A linear mixed model (LMM) was estimated for the ERP and TRF amplitudes, for each subjective workload question, and for the time to complete the surgery task. As the number of mistakes during the surgery task (zero to three mistakes) and the number of times the tissue was damaged provided count data, a generalized linear model (GLM) with a poisson distribution was estimated. The LMM and GLM were estimated using the R package lmer4 (v. 1.1-30). Fixed effects were evaluated using Satterthwaite approximations within the R package lmerTest, which estimates the degrees of freedom to calculate p-values. For the memory task, memory scores were first calculated using edit distance scoring^58^ (i.e., scores range from 0 to 1). Then, the scores received ordinal values which were fit using a cumulative link mixed model using the clmm function of the ordinal package (v. 12-4).

Evidence for an effect was assumed for *α* =.05. We corrected for multiple comparisons using *α* =.05/3 = 0.017 for the ERP/TRF amplitudes (i.e., N1, P2, N2), the comparison between prediction values of the acoustic features (i.e., envelope, noise-reduced envelope, onsets), subjective workload questions (i.e., effort, frustration and distraction), and surgery task performance (i.e., duration, mistakes, and tissue damage).

## Results

For each response we evaluated a condition difference using regression models. We explored whether adding time as a predictor improved the model fit. Table 1 shows the selected model for each response and whether the corresponding beta values were significant.

**Table 1.** The table shows the selected model for each subjective, behavioral, and neural response. ‘Model’ lists the chosen model for each response. Model 1 included ‘condition’ as a predictor and model 2 ‘condition’ and ‘time’ as predictors. Beta values are listed below each predictor. The stars indicate predictors below the Bonferroni-corrected p-value. A trend is marked with a dot.

|  | Response | Model | Condition | Time |
| --- | --- | --- | --- | --- |
| subjective workload questions | tlx-effort | 1 | 3.73* |  |
|  | tlx-frustration | 1 | 2.71* |  |
|  | tlx-distraction | 1 | 1.31* |  |
| behavioral performance | memory score | 1 | -4.63* |  |
|  | duration | 2 | -1.842 | -0.91* |
|  | mistakes | 2 | 0.14 | -0.03* |
|  | tissue damage | 2 | -0.02 | -0.01* |
| ERPs | N1 | 2 | 0.4 | 0.3* |
|  | P2 | 2 | 0.21 | 0.38* |
|  | N2 | 1 | 0.169 |  |
| TRF<sub>envelope</sub> | N1 | 1 | -0.94 |  |
|  | P2 | 1 | -0.37 |  |
|  | N2 | 1 | 0.75 |  |
| TRF<sub>onsets</sub> | N1 | 2 | 0.49 | 0.07 . |
|  | P2 | 1 | 0.025 |  |
|  | N2 | 1 | -0.12 |  |

### Subjective measures of demand

After each block, we asked participants how effortful, frustrating, and distracting they perceived the task. There was a significant increase in scores from the low- to high-demand condition for all three measures (Figure 2 A-C; effort: b=3.73, SE=0.22, p*<*.001; frustration: b=2.71, SE=0.28, p*<*.001; distraction: b=1.31, SE=0.19, p*<*.001). There was no significant change in theses measures over time, i.e., for all three questions the model fit did not improve by adding time as a predictor (supplementary table S2).

**Figure 2.**
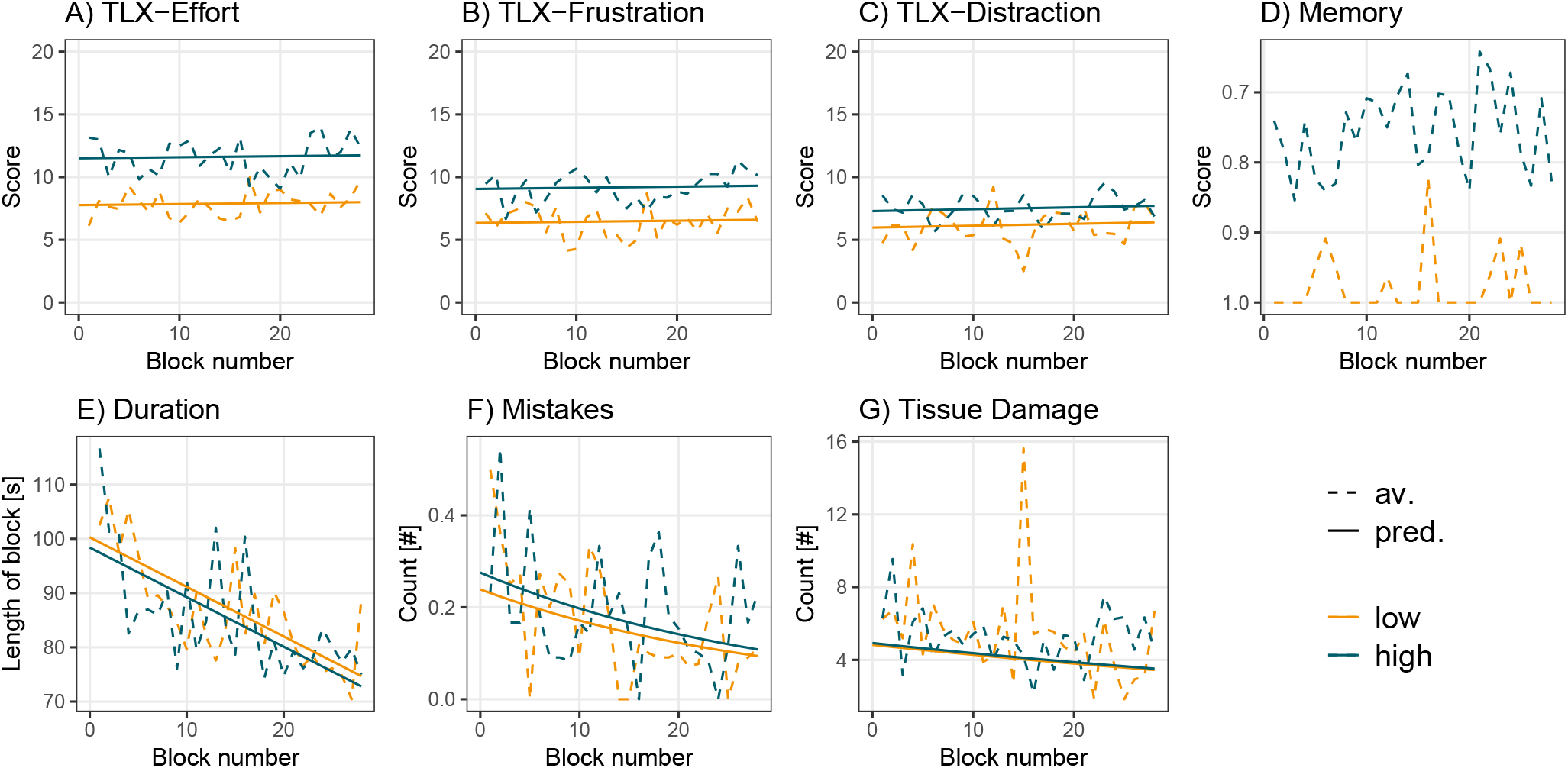
(A-C) Subjective responses. (D) Memory task performance. (E-F) Surgery task performance. Dotted lines show averaged data over participants for each condition and block. Solid lines represent predicted responses that were calculated using the second model. Predicted responses for the memory task were not computed as a commutative link model was used.

### Objective measures of performance

The memory score, which could range between 0 to 1, was in the low-demand condition on average 0.97 (SD=0.13) and in the high-demand condition 0.75 (SD=0.24). The first model, which included only condition as a predictor, was used (supplementary table S2) and revealed a significant decrease in memory score from the low- to high-demand condition (Figure 2D, b=-4.63, SE=0.39, p*<*.001).

To evaluate the performance during the surgery task we used three measures: task duration (i.e., time to finish the surgery task), the number of mistakes (i.e., dropping or rupturing a vessel), and the number of times the tissue was damaged (i.e., tissue that was touched with either of the tools). Figure 2 E-G illustrates the results of the behavioral performance. The performance was already high at the beginning of the experiment, which is evident in the low number of mistakes in most of the blocks. The average number of mistakes is 0.33 (SD=0.48) in the first block and 0.18 (SD=0.39) in the last block. The number of times tissue was damaged changed from 6.2 (SD=3.96) in the first block to 5.4 (SD=4.88) in the last block. Furthermore, task duration appeared to improve, particularly in the first ten blocks, after which it remained relatively stable. For neither of these measures, we found a significant difference between conditions (task duration: b=-1.842, SE=1.57, p=.228; mistakes: b=0.14, SE=0.19, p=.445; tissue damage: b=-0.02, SE=0.034, p=.556). However, for all measures adding time as a predictor significantly improved the model fit (supplementary table S2) and revealed a significant decrease over time (task duration: b=-0.91, SE=0.1, p*<*.001; mistakes: b=-0.03, SE=0.01, p=.0046; tissue damage: b=-0.01, SE=0.02, p*<*.001), i.e. participants became faster and made fewer mistakes.

### Demand and time-on-task effects for ERPs in response to the acoustic letters

The response to the acoustically presented task-irrelevant letters was evaluated by averaging GED component amplitudes for the N1 (Figure 4A), P2 (Figure 4B), and N2 (Supplementary Figure S4A) over their respective time-window. Both the morphology as well as the topographies of the GED components with the strongest eigenvalues (supplementary Figure S3A) are physiologically plausible.

Contrary to our expectation, we observed no significant effect of demand on the N1 (b=0.4, SE=1.84, p=.827), the P2 (b=0.21, SE=1.91, p=.256), or the N2 (b=0.169; SE=2.05; p=.411) amplitude. Exploring time as a predictor significantly improved the models for the N1 and P2 amplitudes but not for the N2 amplitude (supplementary table S3) and revealed a significant N1 amplitude decrease (b=0.3, SE=0.11, p=.0012) and a P2 amplitude increase (b=0.38, SE=0.11, p=.0014) over time.

### Prediction accuracies for the TRFs in response to the OR playback

The TRFs were calculated by relating the continuous OR playback to the EEG signal. We used three acoustic features to calculate the TRF, namely the envelope extracted from the raw audio, the envelope extracted from the noise-reduced audio, and the onsets extracted from the raw audio (Figure 3A). To compare these features, we calculated the prediction accuracy for unseen neural data. Overall, the prediction values were small (Figure 3B) which is common for this measure. Importantly, the envelope of the noise-reduced audio as well as the onsets showed significantly higher prediction accuracies compared to the envelope from the raw audio (noise-reduced envelope: *W* = 3, *p <* .001; onsets: *W* = 1, *p <* .001). The noise-reduced envelope and onsets did not significantly differ from each other (*W* = 118, *p* = .799).

**Figure 3.**
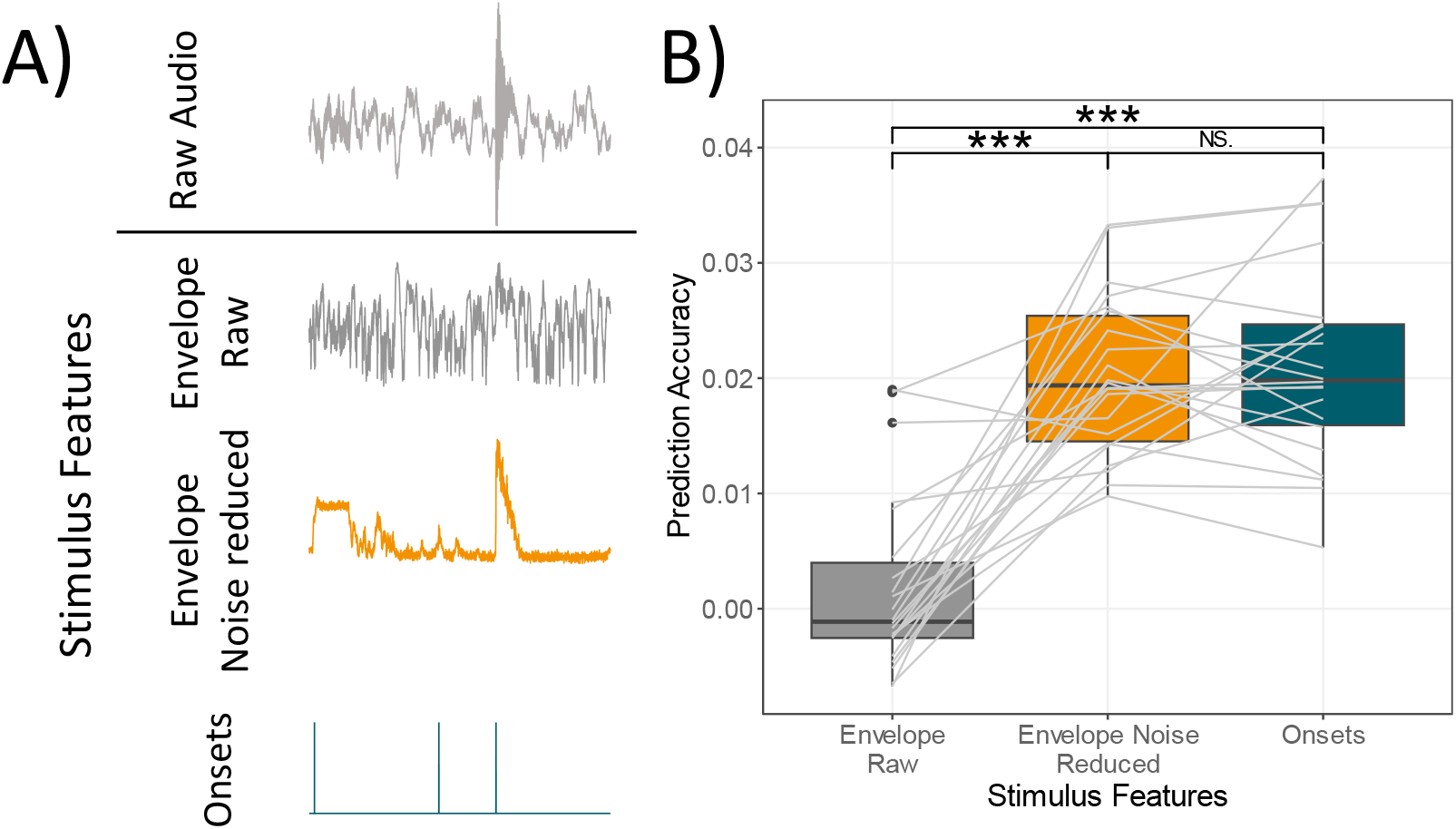
A) The acoustic features used for TRF model estimation. The same stimulus snippet is shown as the raw audio, envelope of the raw audio, envelope of the noise-reduced audio, and onsets of the raw audio. B) Prediction values of each acoustic feature. Each line represents the change in prediction value between acoustic features for each participant. *** p*<*.001

**Figure 4.**
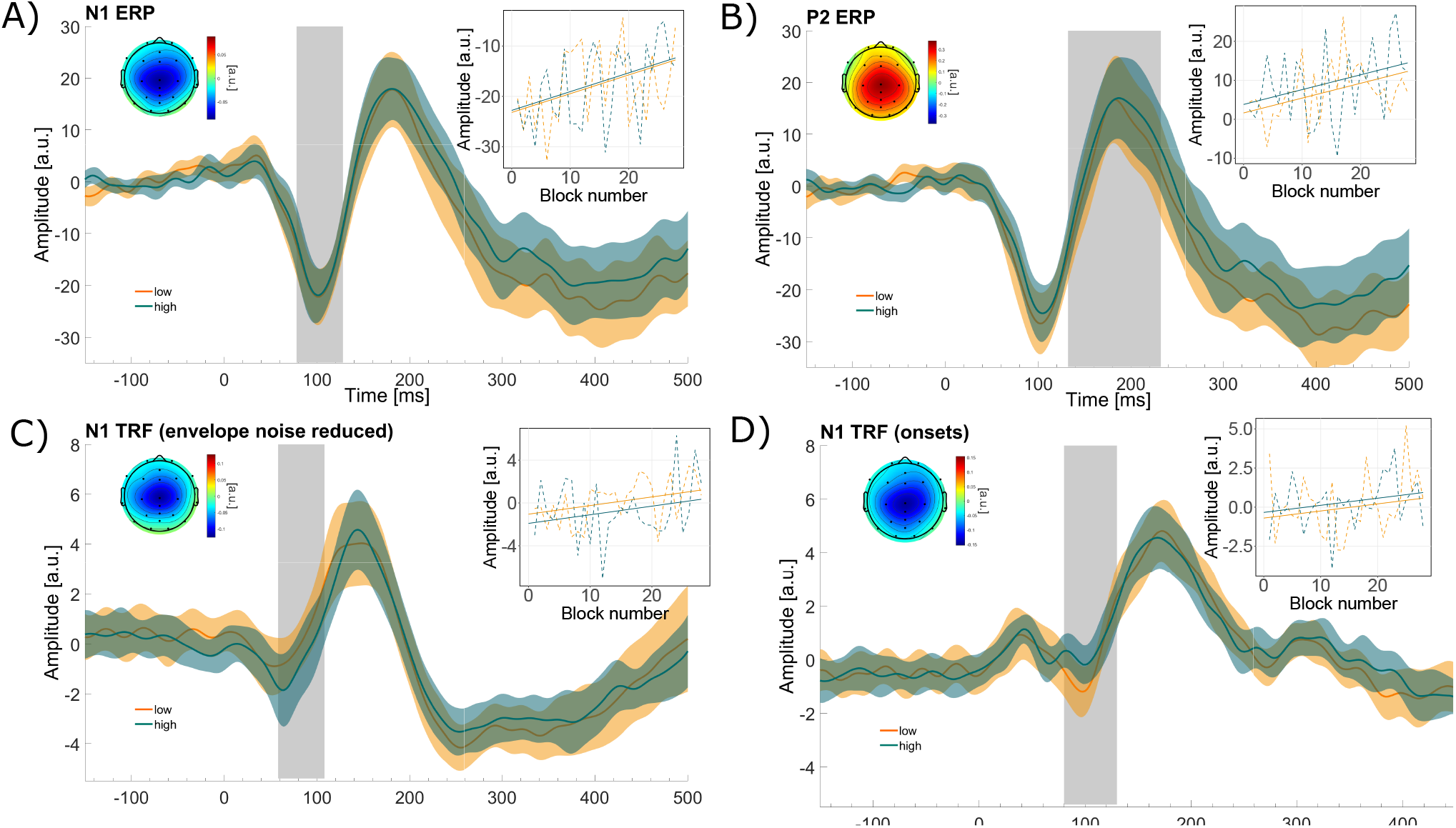
The first GED components of the (A) N1 ERP (demand: n.s., time: p=.0012), (B) P2 ERP (demand: n.s., time: p=.0014), (C) N1 TRF_envelope_ (demand: n.s., time: n.s.), and (D) N1 TRF_onsets_ (demand: n.s., time: n.s.). The morphology shows the GED time-series for each condition, averaged over participants. The shaded area represents the standard error across participants. The grey bar shows the time-window that was used to derive the amplitudes. The topographies depict the forward model of the GED weights and shows which channels contributed most to the GED component. The graph to the right, shows the change in amplitude across time and conditions. Dotted lines show averaged data over participants for each condition and block. Solid lines represent predicted responses that were calculated using the second model.

### Demand and time-on-task effects for TRFs computed from the noise-reduced envelope and onsets

We computed the TRF in response to the noise-reduced envelope (hereafter TRF_envelope_) and in response to the onsets (hereafter TRF_onsets_). Although there was no difference between prediction accuracies between these two acoustic features, they might still represent different aspects of the soundscape and, therefore, different responses. To enhance the signal-to-noise ratio and to spatially filter the data GED components were extracted. The mean amplitudes over the N1, P2, and N2 time-windows were used for evaluation. Figure 4C and D shows the distinct time-series and topographies for the N1 TRF_envelope_ and the N1 TRF_onsets_, respectively. The remaining GED time-series, as well as GED eigenvalues can be found in supplementary figure S3 and S4. Visually there are considerable differences in the temporal evolution between the TRF_envelope_ and the TRF_onsets_. While the TRF_envelope_ follows a trajectory similar to the ERPs its response peaks occur earlier than that of the ERPs and the TRF_onsets_. Regarding the N2 TRF_envelope_, the GED component with the second strongest eigenvalue showed more plausible trajectories and topographies than the first GED component (supplementary figure S4).

As TRF_envelope_ and TRF_onset_ showed similar prediction values, we evaluated amplitudes of both TRFs, and corrected for additional multiple comparisons using *α* =.05/3 (responses) / 2 (acoustic feature) = 0.0083. For all TRF_envelope_ amplitudes, we found no condition difference (N1: b=-0.94, SE=0.57, p=.102; P2: b = -0.37, SE=0.49, p=.449; N2: b = 0.75, SE=0.45, p=.095) and adding time as a predictor did not lead to a better model fit compared to the first model (supplementary table S3).

Regarding the TRF_onsets_ amplitudes we found no condition differences (N1: b=0.49, SE=0.41, p=.23; P2: b = 0.025, SE=0.39, p=.948; P2: b = -0.12, SE=0.35, p=.716). Adding time as a predictor yielded a better model fit for the N1 amplitude, but not for the other amplitudes. This revealed a trend towards significance for the predictor time for the N1 TRF_onsets_ (b=0.07, SE=0.02, p=.011).

## Discussion

In this study, we examined the impact of varying demand and time-on-task on subjective responses, task performance, and EEG responses within a naturalistic setting, utilizing a laparoscopic simulator and an operating room soundscape.

We found divergent subjective and objective behavioral responses. Participants perceived the high-demand condition as more demanding than the low-demand condition, which remained stable over time. However, surgery task performance did not differ between demand conditions but demonstrated improvement over time. The overall experienced demand of the two conditions was evaluated with subjective workload questions. The observed effect that high demand compared to low demand increases subjective workload ratings during a surgery task aligns with prior research^32,34,59^. A possible explanation for the perceived differences in demand across conditions, despite consistent performance levels across conditions, is that participants may have prioritized their performance of the surgery task. By putting more cognitive effort into the surgery task in the high-demand condition, they were able to achieve similar surgery task performance across both conditions. Likewise, a surgeon would prioritize performance during an actual surgery for the benefit of the patient, even if it means an increase in perceived demand. This implies that task performance and subjective experience are not necessarily related and may also explain the contradictory findings reported in the literature. Some studies report a negative impact of increased demand on performance^20,59^, while others show no effect^31^, or suggest that the effect depends on the investigated performance measure^34,60^. Most studies, did not assess the subjective demand, and only concentrate on task performance. However, this limits our understanding how perceived demand and surgical task performance relate and should be addressed in future research by incorporating subjective demand measures.

Regarding the impact of time-on-task on performance, previous research has also demonstrated that task performance improves over time^35,61–63^, highlighting visible training effects within a short amount of practice when using the same surgery task. We did not find a change in perceived demand over time, similar to Suarez et al. (2022)^61^ who found that training did not reduce demand for the same NASA-TLX items as used in this study (i.e., effort and frustration) or the total score. Others reported that surgical training lowers the NASA-TLX total score^35,62^. This once again demonstrates that the relationship between task performance and perceived demand is not straightforward. The use of various methods to vary demand and surgical tasks, each with different performance parameters and subjective and objective measures, also complicates the generalization of findings across studies^64^. To improve generalization, it would be useful to evaluate how the demand experienced during actual surgeries reflects the demand experienced during surgical simulations. Our results suggest that situational demand, rather than actual surgical skill, influences the perceived demand in inexperienced participants. Novice surgeons, who are particularly susceptible to demand^4,31^, may therefore benefit from training in demanding situations to reduce demand during actual surgeries.

As an alternative objective measure to capture the perceived task demand, we investigated two neural response measures to the task-irrelevant background soundscape. Specifically, we analyzed ERPs in response to spoken letters and TRFs in response to the soundscape as a whole. In general, we find a clear neural response to the letters as well as for the naturalistic OR playback, which is in line with our previous study showing significant responses for naturalistic soundscapes^26^. This further expands the use of mobile EEG in applied settings^27^, as both measures can be used to study the neural repose to complex natural soundscape in a work-like environment. Contrary to our hypothesis, we found no significant effect of task demand on the neural measure of sound perception, neither in response to the letters nor to the OR playback. Participants’ ratings indicate that our intended manipulation — to increase the scenario’s challenge — had an effect, with participants perceiving the background sound as more distracting in the high-demand condition. However, this perceptual shift is not reflected in our neural measurements. Our exploratory analyses revealed time-on-task effects, for the N1 and P2, and a trend for the TRF_onsets_ N1. This was apparently unrelated to the subjectively perceived distraction, which remained stable over time.

Our results should be considered in light of the neural measures used to investigate different aspects of the soundscape, i.e., the ERPs and TRFs. The ERPs captured responses to the regularly occurring letters associated with the memory task. To increase the distracting nature of the task-irrelevant soundscape and simulate demanding internal cognitive processes we adapted a serial-recall paradigm. In this study no effect on any ERP time-window was observed, however, previous studies using a serial-recall paradigm have shown that an increase in distraction leads to an increase in N1 ERP amplitude^65^. In contrast to a classic serial-recall paradigm, we used the serial-recall task as a secondary task with a long retention interval (more than 60 seconds), and presented distractors at a rate of 3 seconds. Furthermore, we presented the letters in the presence of background noise, which may have reduced distraction effects in our paradigm^66^. Our choice of distractor presentation (long inter-trial-intervals and in the presence of noise), in combination with an engaging surgery task, may have enabled participants to easily ignore the letters over time, independent of the number of items to recall, resulting in similar responses across conditions. Notably, the results showed a decrease in N1 amplitude and an increase in P2 amplitude, indicating neural adaptation due to repeated stimulus presentation. This may be similar to the adaptation observed for brief repetitive tones^67,68^. The decrease in ERP N1 amplitude could indicate that distraction due to the letters decreased^65^, independent of the conditions. Participants may have initially attended to the letters, e.g., due to their deviation from the continuous soundscape, and learned that the letters are task-irrelevant and can be ignored as such^69^. The P2 amplitude increase is more difficult to interpret, as only few studies investigated it in the context of distraction. One study showed an amplitude increase with increased distraction^70^, which would contradict our interpretation, while another study found no effect of distraction on P2 amplitude^71^. In summary, the results suggest that the task-irrelevant acoustic letters did not sufficiently distract participants to elicit distinct neural responses under varying demands. However, the changes in amplitude across time indicate that the responses can capture some form of modulation. As this was an exploratory finding, further investigation is necessary to reveal how responses to task-irrelevant acoustic events change over time when presented in a realistic work-like setting. This information could shed light on how the brain adapts to such events in real life.

The TRFs captured responses to the realistic OR playback. To the best of our knowledge this is the first study investigating demand effects on responses to task-irrelevant ongoing soundscapes. Despite the fact that we measured a clear TRF to the soundscape, we did not find a difference between conditions. One explanation for this null-finding is that TRFs calculated from the entire soundscape are not sensitive enough to detect demand effects. A small portion of sounds may have elicited demand effects similar to ERPs^39^. However, these effects may have been reduced by mostly demand-unrelated responses. Support for this explanation comes from our previous study^26^, where we found attention-dependent responses for specific highly salient sounds, but only marginal differences in the TRF to the background soundscape. This might also explain why we found no time-on-task effects for the TRF, except for the trend of a decrease for the TRF N1 calculated from the onsets (TRF_onsets_), despite the effect for the ERPs. While the ERPs captured responses to similar sounds, the TRFs captured responses to the OR-playback which was much more diverse than the letters. This might have decreased a potential time-on-task effect for the TRFs compared to ERPs, as different and overlapping responses are less sensitive to detect effects compared to similar and isolated responses^42^. For example, an audio signal, that contains a constant high level of noise, would result in an essentially flat envelope, for which no reliable TRF could be computed. Any acoustic event, that is embedded in this background sound, though perceivable to the human ear, would not be reflected in the envelope. This is evident in our finding that TRFs computed from the raw envelope, which captured little variation of the soundscape, provided low prediction values. The individual acoustic events, such as the beeps of machinery or the clatter of tools, were only adequately captured by reducing the noise in the raw envelope. By highlighting the individual acoustic events, the envelope became more similar to the computation of the onsets. The similarity between these two features is expressed in similar prediction values. This suggests that the brain’s processing of sounds goes beyond simply following the acoustic envelope^41,72^. Instead, it actively distinguishes and monitors individual acoustic events in the presence of background noise, similar to how speech is perceived in noisy environments^73^. Note, that similar prediction values, do not necessarily imply a similar neural representation of the soundscape^42^. This is hinted by the trend in the TRF_onsets_ N1 which was not observed in the TRF_envelope_. However, the similar prediction accuracies indicate that onset information about acoustic events is enough to estimate the neural response to complex soundscapes. This can have practical implications when privacy is important (e.g. in an actual OR context where medical information about a person are discussed). It would be sufficient to extract acoustic onsets automatically, without the need to record the raw audio^40,74^. Overall, we have demonstrated that it is possible to obtain responses from continuous task-irrelevant soundscapes. However, future studies may benefit from describing the soundscape not only in the form of an envelope or by extracting onsets, but to differentiate between different sound sources to obtain a more fine grained picture.

An alternative explanation for the discrepancy between our objective measures (i.e., task performance and neural measures) and the subjective reports is that the subjective reports did not accurately reflect participants’ perceived demand throughout an entire block. As the subjective responses were collected at the end of a block, they may reflect the perceived demand of the memory task during encoding. In other words, participants found it difficult to remember two compared to eight letters, but experienced the surgery task as similar demanding across conditions. Given the complexity of the task, accurately pinpointing the source of demand — whether it’s remembering the letters, the surgery task, or a combination of both — might be difficult for participants. Indeed, the NASA-TLX has been criticised that different participants might rate different aspects of the task^75^. Collecting separate subjective responses for each task may have improved interpretation. However, in natural environments, the source of demand, whether it is the environment or the task, can rapidly change, making subjective responses at a single point in time difficult to interpret. This once again emphasizes the importance of continuously and objectively monitoring sources of demands, such as the soundscape, to understand how they affect the individual.

In order to improve the understanding of varying demands on responses to ongoing soundscapes, we offer two suggestions for future work. Firstly, a different manipulation of demand may provide further insight into the feasibility of obtaining demand-dependent responses to naturalistic soundscapes. We intentionally used a demand manipulation that did not require an overt response so that participants could focus on the surgery task, a scenario we believe is more realistic than asking the surgeon to constantly interrupt the surgery to press a response button. The disadvantage of this approach is that we did not observe a demand-related effect on our objective measures and therefore do not know how effective the demand manipulation was. An alternative way to manipulate demand could involve surgery tasks of varying difficulty levels, which would provide a realistic scenario. However, this approach may hinder the comparison of performance measures. Secondly, we may have to improve our neural measures. A more detailed description of the soundscape could enhance TRF calculation. Instead of using only the envelope of the soundscape, as we have done here, a more detailed description of the soundscape, e.g. by differentiating between specific sounds may lead to a better estimate of the neural response^42^ and hence may allow us to detect more subtle difference between the conditions. For example, individual sounds can be evaluated based on their salience^76^, which can help assessing their potential to distract. These suggestions could improve the use of EEG in work environments and enhance our understanding of whether and how cognitive states can be inferred from responses to natural soundscapes.

Our ultimate goal is to study the neural response to complex soundscapes in everyday life. In order to learn from our results for other studies that are interested in everyday soundscapes it is important to discuss some properties of the soundscape we have used here. We used the playback of a natural soundscape that was recorded in an OR using a static microphone positioned at the center of the room. We presented this soundscape via two loudspeakers to a stationary participant. This setup ensured that the soundscape maintained realistic spatial properties from the listener’s perspective, reflecting the acoustic environment of an operating room. Despite the naturalness of this soundscape, there are important differences between the OR playback we used and an actual OR soundscape experienced in real-life situations. One key difference is sound expectancy. For instance, in an OR setting, there is an expectation that specific actions one can see, such as laying down a tool, will produce a particular sound. In other words, there is a congruence between visual and auditory information. In our study, this audio-visual congruence is absent; all the sounds our participants hear are not congruent with the actual situation they are in. One might expect these differences to affect the neural response we have measured, although the direction of this effect is difficult to predict. Some speech tracking studies have shown an enhanced tracking of the speech envelope when congruent visual information is provided^77^. On the other hand, a mismatch of visual and auditory information could also lead to a larger neural response^78^. A second difference between the soundscape experienced in the OR and our OR playback is the relevance of the sounds for the participant. In an OR setting, many sounds carry contextual information — be it the use of particular tools or the request for instruments — that can direct attention or elicit specific task-related actions; some sounds, however, can be safely ignored by the surgeons as they do not carry any meaning. In contrast, the soundscape we presented was entirely task-irrelevant for our participants, essentially transforming all sounds into a meaningless noise background. One can expect that for personally relevant or salient information within a soundscape influences its processing^25,79,80^. Taken together, these factors may significantly change the neural response to the naturalistic soundscape we have presented, compared to when this naturalistic soundscape would have been experienced in its actual context. Moving forward, future studies should be aware of these important factors when studying naturalistic soundscapes.

Our research represents a significant advancement in the application of EEG to provide reliable insights into brain responses within contexts that closely resemble real workplace environments. We included a surgical simulation involving bi-manual tasks, visual feedback, and varying cognitive demands amidst a naturalistic soundscape. This demonstrated the feasibility and potential of combining EEG and audio recordings to assess cognitive processes in complex environments. The study emphasizes the challenge of identifying neural markers that align with subjective experiences while maintaining ecological validity and experimental control. By addressing these challenges, our research not only contributes to the methodological advancements in neurophysiological studies in real-world settings but also opens avenues for enhancing surgical performance through improved understanding and optimization of the auditory environment.

## Data availability

The datasets generated during and/or analyzed during the current study are available from the corresponding author on reasonable request.

## Acknowledgements

This work was funded by the Deutsche Forschungsgemeinschaft (DFG, German Research Foundation) ID 411333557 & 490839860, and by the Forschungspool Funding of the Oldenburg School of Medicine and Health Science. We thank Agnes Hanssen for her help in data acquisition.

## Author contributions statement

M.R. and M.B. conceptualized the experiment. M.R. performed the data acquisition and analyzed the data, which was assisted by T.H., M.J., and M.B.. M.R. wrote the manuscript to which T.H., M.J., V.U., and MB contributed with critical revisions. All authors reviewed the manuscript.

## Additional information

The author(s) declare no competing interests. During the preparation of this work the authors used ChatGPT (v. 3.5) and DeepL Write (acadamic style) to improve grammar and wording. After using these tools, the authors reviewed and edited the content as needed and take full responsibility for the content of the publication.

