## Supplementary Material for "Sound perception in realistic surgery scenarios: Towards EEG-based auditory work strain measures for medical personnel"

### Deviations from preregistration

#### TRF analyses

- Initially we ran the cross-validation procedure 100 times and randomly chose 90% of the blocks as test data and the remaining 10% of the blocks as training data. However, we realized that this is computationally expensive while resulting in very similar prediction values as with the approach in the manuscript.
- During the last step of TRF calculation we computed the window from -220 to 500 ms. In the preregistration the window ranged from -200 to 500 ms, however, due to an early artefact common to TRF calculations[54] we had to increase the window.
- Due to an oversight we did not detail in the preregistration how we determined the individual optimal lambda value.
- We z-standardized the TRF values before GED analyses. This was necessary as the individual lambda range was large, thus resulting in individual TRFs on different scales which lead to violations of the normality assumption for the statistical comparison.

**GED analyses** Our initial idea was to individualize the analyzes pipeline, especially the GED computation and compute a spatial filter per participant. However, it turned out that this worked well for our pilot data but to a much lesser degree for the entire dataset. The GED analyses led to component time-series and maps that were hard to interpret or not meaningful making component selection difficult. Therefore, we used a grand average GED model and made the following adaptations:

- We used the GED weights calculated from all subjects and applied them to each subject and block. The initial idea was to calculate GED weights for each participant calculated from the grand average over blocks and apply the weights to all blocks of of this participant.
- We calculated fixed time-windows to determine the component time-windows. Initially, the full width at half maxmimum/minimum with respect to the peak was used, but this lead to time-windows that extended over the response of interest.
- We calculated the S and R covariance matrix for each participant and cleaned the matrix before averaging. This is a step included in Cohen (2022)[57]. Initially, we used the grand average ERP of a participant to compute one S and one R matrix, thereby making the cleaning step unnecessary.

Further adaptions to the GED analyses were made:

- Initially the peak of all ERP and TRF components were searched for in the same time-window. However, it turned out that the envelope response peaks earlier than the ERPs and TRFs calculated from the onsets due to temporal smearing. To account for this, we searched for the envelope peaks in an earlier time-window, as stated in the manuscript.
- In the preregistration the lambda value to regularize the R matrix was defined as $\lambda = 0.1$ which was a typo. The initial analyses was set up with $\lambda = 0.01$ as stated in the manuscript and which was also used in Cohen (2022)[57].

#### Statistical analyse

- The initial idea was to calculate the maximum LMM (i.e., $\hat{y} \sim condition + (condition|participant)$). However, in most cases this model did not converge, especially when adding time as a predictor. We decided to start with model 1, then successively add fixed effects, and finally evaluate the change in model fit.
- For the memory scores, we initially wanted to calculate a beta regression model, but realized that these are not applicable when responses are bound to 0 or 1, which was the case for our data.

### Shift of letter onset

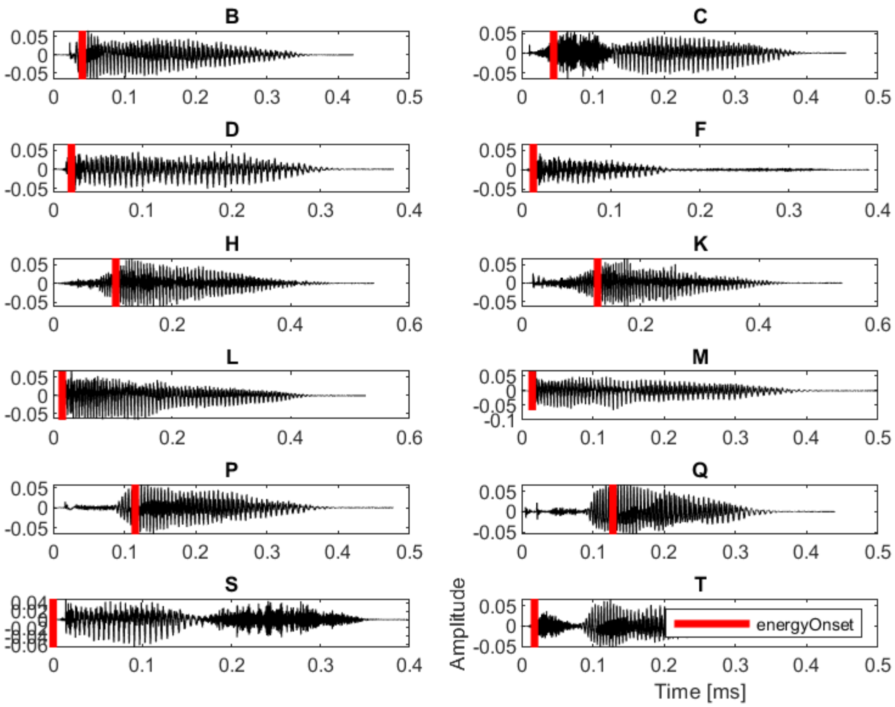


**Figure S 1.** Time-representation of each letter. The red line indiactes the shift of the letter onset.

### Removed IC containing muscle movement

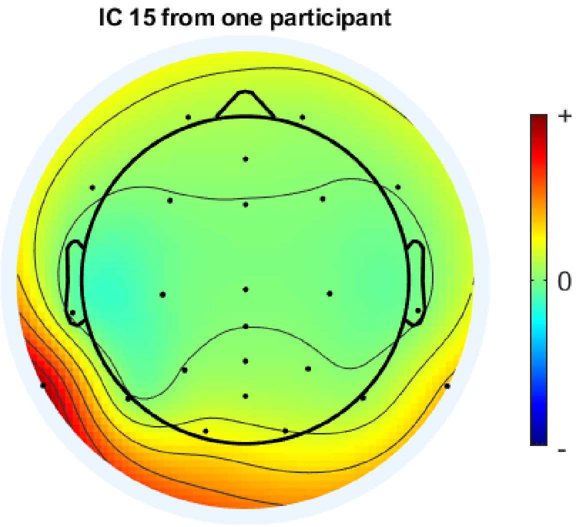


**Figure S 2.** This is an example of an independent component of one participant that was manually removed, as it contained noise at the mastoid electrode.

### GED covariance matrices

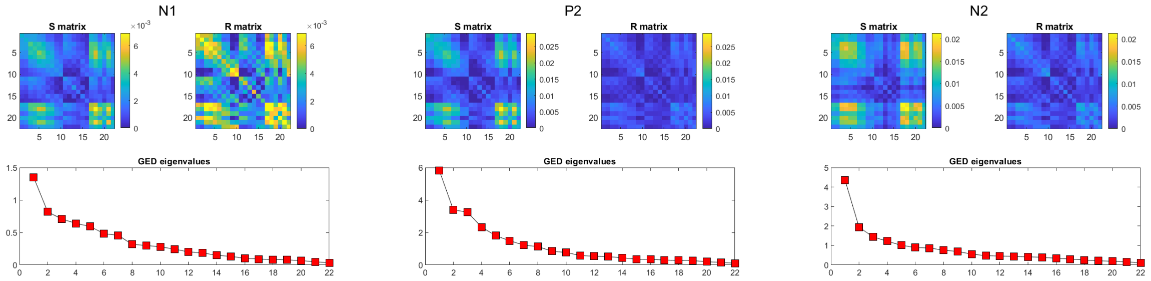


**(a)** ERP components in response to the letters.

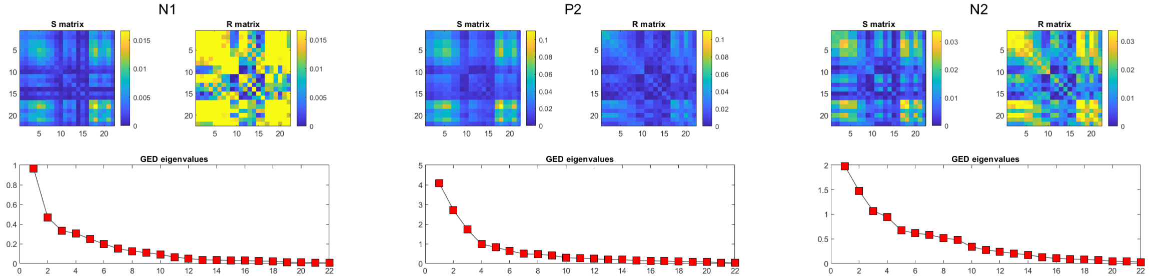


**(b)** TRF components calculated from the noise reduced envelope.

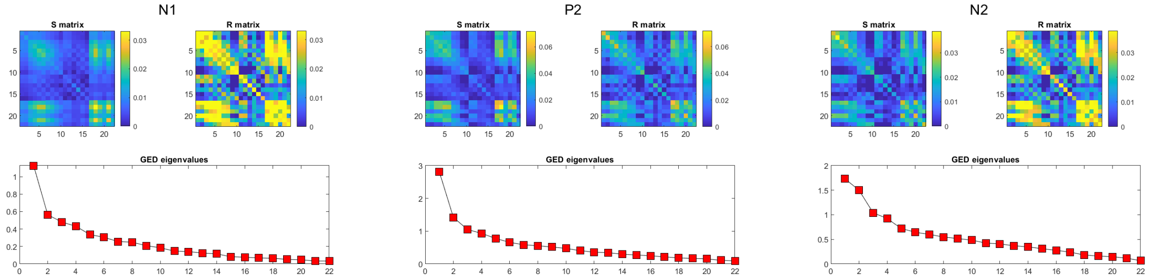


**(c)** TRF components calculated from the noise reduced envelope.

**Figure S 3.** GED covariance matrices and eigenvalues for (A) ERPs, and TRFs calculated from the (B) noise reduced envelope and (C) onsets. Upper left and right plot depict the $\bar{S}$ and $\bar{R}$ covariance matrix, respectively. The lower plot depicts the eigenvalues of each component.

### GED components

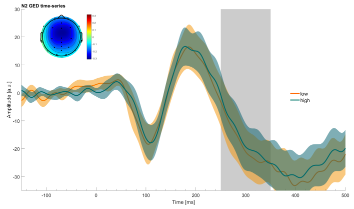


**(a)** ERP N2: First GED component.

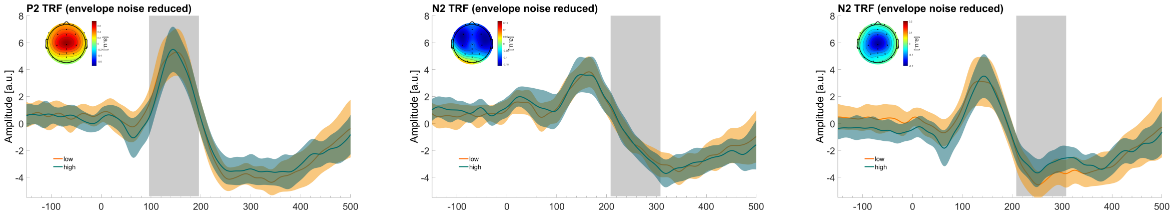


**(b)** **(left)** First GED component of P2 TRF envelope (noise-reduced). **(middle)** First GED component of N2 TRF envelope (noise-reduced) and **(right)** second GED component of N2 TRF envelope (noise-reduced). The second component was chose for the N2 analyses, as the map and topographies look more plausible.

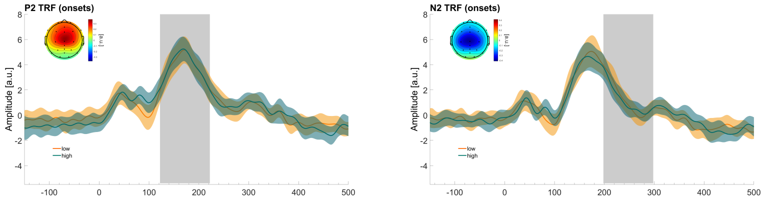


**(c)** **(left)** First GED component of P2 TRF onset. **(middle)** First GED component of N2 TRF onset.

**Figure S 4.** This figure shows GED components that were analyzed but not shown in the manuscript. The GED time-course displays averaged data (line) and standard error (shaded area) across participants. The grey area shows the time-window that was used to compute the average amplitude. The topographies depict the forward model of the GED component.

### Statistical model compairsons

#### *Behavioral and subjective Results*

| response | predictor | AIC | BIC | logLik | Chisq | Df | Pr(>Chisq) |
|---|---|---|---|---|---|---|---|
| tlx-effort | condition | 3164.4 | 3182.1 | -1578.2 | | | |
| | condition+time | 3166.0 | 3188.2 | -1578.0 | 0.3593 | 1 | 0.5489 |
| | condition*time | 3166.7 | 3193.3 | -1577.3 | 1.3156 | 1 | 0.2514 |
| tlx-effort | condition | 3440.7 | 3458.5 | -1716.4 | | | |
| | condition+time | 3442.5 | 3464.7 | -1716.2 | 0.2661 | 1 | 0.6059 |
| | condition*time | 3444.4 | 3471.0 | -1716.2 | 0.0961 | 1 | 0.7565 |
| tlx-distraction | condition | 3006.0 | 3023.7 | -1499.0 | | | |
| | condition+time | 3006.4 | 3028.7 | -1498.2 | 1.5193 | 1 | 0.2177 |
| | condition*time | 3007.8 | 3034.5 | -1497.9 | 0.6005 | 1 | 0.4384 |
| Performance time | condition | 5656.2 | 5674.0 | -2824.1 | | | |
| | condition+time | 5576.4 | 5598.6 | -2783.2 | 81.8695 | 1 | <2e-16 *** |
| | condition*time | 5577.7 | 5604.3 | -2782.8 | 0.6821 | 1 | 0.4089 |
| Mistakes | condition | 619.97 | 633.29 | -306.98 | | | |
| | condition+time | 613.99 | 631.74 | -302.99 | 7.9818 | 1 | 0.004725 ** |
| | condition*time | 614.66 | 636.86 | -302.33 | 1.3249 | 1 | 0.249721 |
| Tissue damage | condition | 3807.9 | 3821.2 | -1901.0 | | | |
| | condition+time | 3778.8 | 3796.6 | -1885.4 | 31.0797 | 1 | 2.476e-08 *** |
| | condition*time | 3778.8 | 3801.0 | -1884.4 | 1.9995 | 1 | 0.1573 |

**Table S 2.** The table shows model performance of the statistical models. The simplest model contained condition as a predictor. There is no model comparison for the memory condition, as the models did not converge when adding time as a predictor. Signif. codes: 0 ‘***’ 0.001 ‘**’ 0.01 ‘*’ 0.05 ‘.’ 0.1 ‘ ’ 1

***EEG data***

| response | predictor | AIC | BIC | logLik | Chisq | Df | Pr(>Chisq) |
|---|---|---|---|---|---|---|---|
| ERP - N1 | condition | 5484.0 | 5501.5 | -2738.0 | | | |
| | condition+time | 5475.5 | 5497.4 | -2732.7 | 10.4946 | 1 | 0.001197 ** |
| | condition*time | 5476.2 | 5502.6 | -2732.1 | 1.2575 | 1 | 0.262118 |
| ERP - P2 | condition | 5549.9 5567.5 | -2770.9 | 5541.9 | | | |
| | condition+time | 5541.7 | 5563.7 | -2765.9 | 10.1876 | 1 | 0.001414 ** |
| | condition*time | 5543.3 | 5569.6 | -2765.6 | 0.4539 | 1 | 0.500484 |
| ERP - N2 | condition | 5622.4 | 5639.9 | -2807.2 | | | |
| | condition+time | 5623.6 | 5645.6 | -2806.8 | 0.7320 | 1 | 0.3922 |
| | condition*time | 5625.5 | 5651.9 | -2806.8 | 0.0959 | 1 | 0.7568 |
| TRFenv - N1 | condition | 4067.9 | 4085.5 | -2030 | | | |
| | condition+time | 4066.6 | 4088.6 | -2028.3 | 3.2459 | 1 | 0.0716 . |
| | condition*time | 4068.2 | 4094.6 | -2028.1 | 0.4148 | 1 | 0.5196 |
| TRFenv - P2 | condition | 3887.3 | 3904.9 | -1939.4 | | | |
| | condition+time | 3889.2 | 3911.2 | -1939.6 | 0.0451 | 1 | 0.8318 |
| | condition*time | 3891.2 | 3917.6 | -1938.6 | 0.0131 | 1 | 0.9090 |
| TRFenv - N2 | condition | 3778.3 | 3795.9 | -1885.2 | | | |
| | condition+time | 3780.3 | 3802.3 | -1885.1 | 0.0175 | 1 | 0.8946 |
| | condition*time | 3782.3 | 3808.6 | -1885.1 | 0.0258 | 1 | 0.8723 |
| TRFons - N1 | condition | 3663.6 | 3681.2 | -1827.8 | | | |
| | condition+time | 3659.2 | 3681.2 | -1824.6 | 6.3816 | 1 | 0.01153 * |
| | condition*time | 3661.2 | 3681.5 | -1824.6 | 0.0408 | 1 | 0.83994 |
| TRFons - P2 | condition | 3610.4 | 3628.0 | -1801.2 | | | |
| | condition+time | 3610.1 | 3632.1 | -1800.1 | 2.2199 | 1 | 0.1362 |
| | condition*time | 3610.6 | 3637.0 | 1799.3 | 1.5038 | 1 | 0.2201 |
| TRFons - N2 | condition | 3440.1 | 3457.7 | -1716.0 | | | |
| | condition+time | 3439.8 | 3461.8 | -1714.9 | 2.2371 | 1 | 0.1347 |
| | condition*time | 3441.8 | 3468.2 | -1714.9 | 0.0232 | 1 | 0.8790 |

**Table S 3.** The table shows model performance of the statistical models. The simplest model contained condition as a predictor. TRFenv: TRFs were calculated using the noise reduced envelope. TRFons: TRFs were calculated using the onsets of the raw audio. Signif. codes: 0 ‘***’ 0.001 ‘**’ 0.01 ‘*’ 0.05 ‘.’ 0.1 ‘ ’ 1
